# Principal Component Analysis applied directly to Sequence Matrix

**DOI:** 10.1101/336115

**Authors:** Tomokazu Konishi, Shiori Matsukuma, Hayami Fuji, Daiki Nakamura, Nozomi Satou, Kunihiko Okano

**Affiliations:** Department of Biological Environment, Faculty of Biresource Sciences, Akita Prefectural University, Shimoshinjyo Nakano, Akita, 010-0195 Japan; Biodiversity Promotion Division, Tokyo Branch, Regional Environmental Planning Inc. NDS Building 2-22-3 Sakurashinmachi Setagaya-ku, Tokyo, 154-0015 Japan

**Author notes:** Corresponding author: Tomokazu Konishi.

**Keywords:** phylogeny, evolution, horizontal gene transfer, dendrogram, network, cladistics, mutation, falsifiability, objectivity, mathematical model

## Abstract

Sequence data is now widely used to observe relationships among organisms. However, understanding structure of the qualitative data is challenging. Conventionally, the relationships are analyzed using a dendrogram that estimates a tree shape. This approach has difficulty in verifying the appropriateness of the tree shape; rather, horizontal gene transfers and mating can make the shape of the relationship as networks. As a connection-free approach, principal component analysis (PCA) is used to summarize the distance matrix, which records distances between each combination of samples. However, this approach is limited regarding the treatment of information of sequence motifs; distances caused by different motifs are mixed up. This hides clues to figure out how the samples are different. As any bases may change independently, sequence is multivariate data essentially. Hence, differences among samples and bases that contribute to the difference should be observed coincidentally. To archive this, the sequence matrix is transferred to boolean vector and directly analyzed by using PCA. The effects are confirmed in diversity of Asiatic lion and human as well as environmental DNA. Resolution of samples and robustness of calculation is improved. Relationship of a direction of difference and causative nucleotides has become obvious at a glance.

## Introduction

Nucleotide sequence could be desired information for classification of organisms such as performed in phylogenetics, as genetic information is highly specific to individuals, easy to obtain with accuracy, and may reflect biological characters of samples. Indeed, amplifying specific fragments of DNA and obtaining nucleotide sequences have become ubiquitous tool for this purpose [1,2]. However, there is no simple solution to estimate relationships among the sequences. As they are qualitative data, numerical conversion is required for any calculation for estimating the relationships in a quantitative way. Additionally, a nucleotide sequence is a multivariate data with huge number of independent items that are recorded as form of bases. Therefore, differences among samples consists of multiple dimensions, and are difficult to be understood. In this sense, estimation of sample relationships is a question in multivariate analysis in essence.

Conventionally, relationships among nucleotide sequences are summarized using a dendrogram. Two classes of approaches are available to estimate the dendrogram (Fig 1). One targets the distance matrix, which records distances among samples, and both the estimation of the distances and the manner in which they are summarized to make the dendrogram are based on specific mathematical models. The other targets directly each of the sites (a set of nucleotides or amino acids in a same position of aligned sequences) and calculates the score for a certain tree model; the estimation of the score is based on a given model that assumes several parameters. (Fig 1, maximum parsimony, likelihood and Bayesian). Via heavy computational calculations, optimized values are found for the parameters (basis of the calculations are explained in detail in a textbook of Yang (2006); moreover, please refer to the on-target short review by Yang & Rannala (2012).

**Fig 1.**
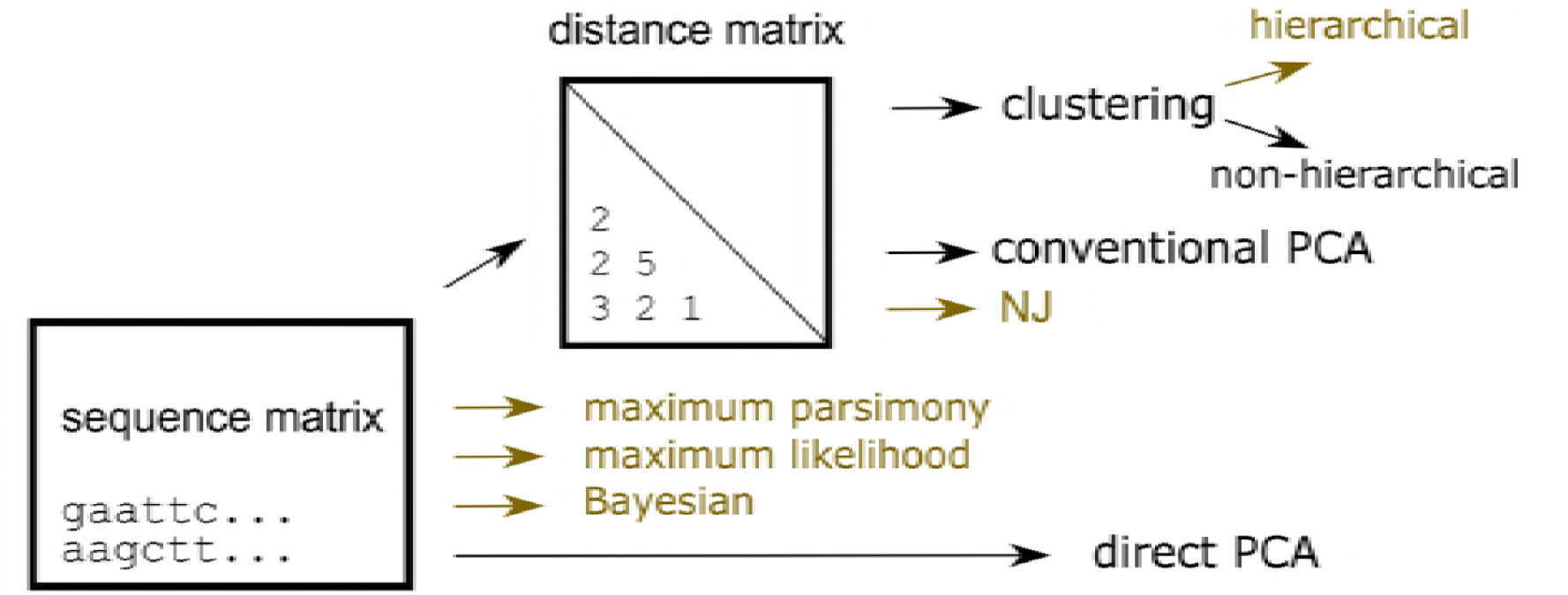
Schematic diagram of the methods used in the study. Two distinct approaches are used: one estimates the distances among samples, to form a distance matrix using one of various models (Yang, 2006), and then finds the relationships based on the distances. The other finds the relationship directly from the sequence matrix. Only direct PCA shows the relationships of samples and bases coincidently. Methods that estimate a tree-shaped relationship are colored in oak.

Despite the efforts to find better dendrograms, the results essentially lack objectivity. Although the mathematical models are designed taking evolution into consideration, they are based on many assumptions that could never be verified by evidence. In fact, most individuals that participate in the process of evolution have been lost, and evidence of periods of time are only rarely available. The topology of a tree, i.e., which combination of samples should be connected, belongs to models, and not to parameters; such assumptions of the tree shape can generate errors, referred to as *long-branch attraction* [3], that are difficult to identify. Moreover, knowledge obtained from such dendrograms cannot be integrated beyond different models. Accordingly, the resulting dendrograms are based on certain ideas that are not falsifiable; in this sense, their standpoint does not fall within conventional science, as is the case of superstring theory [4].

Mathematical models often assume several conditions. For example, those that produce a phylogenic tree assume that all the subjected samples share a tree-shaped relationship (in Fig 1, Methods, colored in oak). In general, such universal propositions can be disproved by showing an opposing example. Below is a simple example of the tree-shape assumption.

In speciation, it was long believed that accumulating mutations generate new species; this makes a linkage between the new and ancestral species, forming a branch of a phylogenic tree. However, whole genome sequencings of prokaryotic species have revealed that evolution is not limited to such hierarchical way, but genes may be horizontally transferred among species [5]. Indeed, such transfer may occur among eukaryotes [6]; additionally, remixing of genetical information often occurs by sexual mating. Such transfer connects distant branches, and hence the relationships among organisms would be a complex network, not a tree. As Neighbor Joining and other clustering method summarize the relationships to estimate phylogenic tree, if the real structure was a network, the resulted estimation will be distorted. It is obvious that false assumptions should not be applied to scientific studies.

In the classification of samples, other non-hierarchical methods have been used, in addition to the tree-based approach (Fig 1). A typical example is k-means; for the given number of clusters, this absorbs neighbor samples of randomly specified starting samples. The self-organizing map is another example [7]; this is an artificial neural network that organizes samples with low-dimensional nodes, which is preferentially designated by the analyst. The two approaches basically do not indicate the distances among samples. Moreover, the results are heavily dependent on the given numbers or dimensions, and reproducibility of the results cannot be expected.

Scientific data analyses must fulfill several specific requirements, such as objectivity and reproducibility [8]. Science is an attempt at integrating the contributions of many researchers. Knowledge without objectivity or reproducibility ruins the integration process. The approaches shown above are not objective, because their assumptions are never falsified by the real data. Moreover, they are not reproducible, because many arbitrary options can drastically change the results. Of course, it is erroneous to keep using inappropriate models, because they would lead to the wrong conclusions.

It is not easy to estimate a phylogenic network, and an appropriate answer would not be provided by an automated method. Short DNA sequences are insufficient to estimate relationships among organisms; the result may be false, and not falsifiable. Such an attempt does not constitute science, but a gamble, regarding objectivity; rather, we need to be aware of the limitations of the existing information. However, as the data structure could be complex, i.e., the base sequence may have the same dimension as its length, a mathematical aid for observing the structure of data is required. A method that can observe the sequence matrix data without harming its structure is preferred; if we can preserve the structure, we would be able to integrate the results to estimate the network structure when sufficient information becomes available.

To maintain objectivity, we should not connect samples based on unverifiable assumptions; rather, they should be kept pendulous. Analyzing the distance matrix using Principal Component Analysis (PCA) would satisfy this criterion because it does not assume a specific structure of data (Fig 1, conventional PCA). Rather, it rotates the matrix and projects it to sets of diagonal axes; it finds directions of differences and presents them by using the axes. The relationships are presented within the found axes with a simpler vision than the original matrix. Both the reproducibility and objectivity of the calculations are satisfactory. This method seems to be used rather spontaneously and spread, as PCA is a preferred method for analyzing matrix data such as the distance matrix. An early example was in a textbook [9].

However, as this method targets the distance matrix, all information for the sites is lost and is not included in the results; i.e., which site contributes to which difference and how it occurs. This is a serious disadvantage in the estimation of the role of sequence motifs or finer relationships between samples. As each sample of sequence data are made of multiple bases, they can be recognized as a point located in multiple dimensions of the same number of bases. Difference between a pair of samples defines a vector in the dimensions, which are consists of both length and direction. Direction could be important as well as distance, therefore, it is better to analyze information of bases and samples coincidently.

To archive this, sequence data have to be transferred to numeral. Here they are replaced as boolean vector that is presented by 0 and 1. For example, a base of DNA is recorded by a set of five digits (A, T, G, C, and -); in this notation, T can be expressed as “01000”. This transformation has merits that no information is lost and so it is completely reversible; also, it is applicable to both nucleotide and amino acid sequences. Differences between two samples are defined by subtraction, distance is found by using Euclidean length, and average sequence is found as the arithmetic mean of samples. Some of the options for adjustments for weights, parallel, convergent, or back substitutions may be still possible, as will be discussed later. Then the quantified sequence matrix is analyzed by PCA (Fig 1, direct PCA).

The PCA applied here (Konishi, 2015) was based on Euclidean metric space; hence, it is compatible with randomly occurring mutations or a Brownian model. From this point of view, it is completely different from several methods proposed for phylogenetics, such as size-correction or phylogenic PCA (Revell, 2009; Uyeda, Caetano, & Pennell, 2015). Those methods were developed to fit non-Brownian evolution; however, they had to assume many conditions that are never verified through analysis. Hence, they are outside the scope of this article, which introduces a scientific method.

The direct PCA method introduced here analyzes a sequence matrix by PCA. Alexe et al. analyzed a Boolean matrix by PCA, not a distance matrix [10]; in this method, the Boolean displayed different bases from a consensus sequence. Hence, it analyzes positions of base alterations, but information pertaining to sequence motifs is negated. Thus, it is different from any of those shown in Fig 1. In direct PCA, both PC for samples and bases are calculated. The latter are loadings given the full length; they indicate which nucleotide base (or amino acid residue) contributes to the scores of the former, and vice versa. By normalizing for the size of the matrix, they show absolute values that can be compared among studies [11]. According to the nature of the PCA, the structure of the data is maintained, whereas other methods distort the structure by applying certain estimations.

Here the method was applied to observe divergence in major histocompatibility complex (MHC) gene and to classification of environmental DNA. Each axis of PCA presented differences of samples and bases coincidentally: which base or motif was characteristic to which group of samples and how. This helped understanding differences among multiple samples. The directivity of each axis improved robustness to outlying bases, which appear rarely among samples, and improved resolution of samples when applied to phylogenetic purposes.

Robustness is an important characteristic when evaluating analytical methods. In some cases, arbitrary parameters used in analyses could change the global results; for example, gap penalty is a parameter of alignment, and can change the number of gaps that are inserted in the aligned sequences. If this parameter affects the results of the analysis significantly, the analysis lacks objectivity, because the analyst may control the results by choosing the parameter. Moreover, outlying samples would also change the results; by adding or removing the outliers, analysts may control the results. Therefore, robustness could be comprehended as a concept that indicates the resistivity of a method to such arbitrary choices.

## Results

### Structure and presentation of data

First, let us examine an example that may indicate a characteristic of the network structure. In Fig 2A, relationships among sewage bacteria taken from active sludge were estimated with some reference species by using distance matrix and NJ method. This method finds the tree structure from distance matrix of the sequences, estimating minimum evolution to connect the whole samples. The resulted tree shows several clusters connected rather complex ways, and some of them are apart from reference species. Of course, the contributions of sequence motifs do not appear in the presentation.

**Fig 2.**
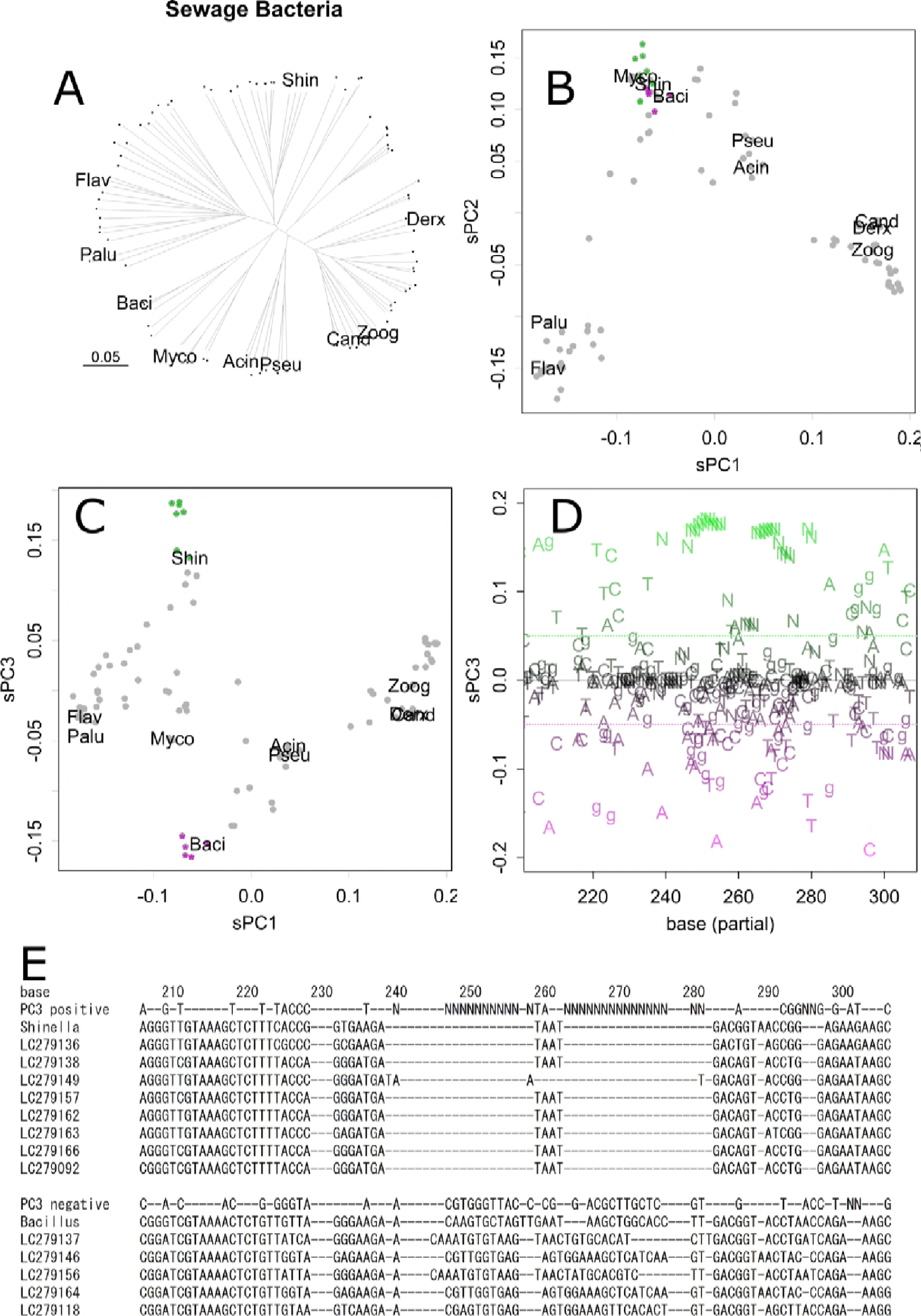
Analysis of sewage bacteria. Neighbor Joining tree. Each dot presents a sample. All the samples are connected to the tree by a single line. Abbreviations are the first four letters of reference samples for *Acinetobacter indicus, Bacillus anthracis, Candidatus Accumulibacter phosphatis, Derxia gummosa, Flavobacterium pectinovorum, Mycobacterium sp. strain FED_F02A, Paludibacter propionicigenes, Pseudomonas punonensis, Shinella kummerowiae,* and *Zoogloea resiniphila,* respectively (**A**). Scatter plot of PC1 and PC2 for samples (**B**). Scatter plot for PC1 and PC3 (**C**). PC3 for nucleotides (partial) (**D**). Positive and negative bases are colored green and magenta, respectively; those had low contributions are colored gray, and none contribution are omitted. *Shinella*, which appeared as a positive in the same PC3 axis (**C**), tended to contain bases indicated in green, whereas *Bacillus* tended to contain bases indicated in magenta. Such tendencies are obvious in a summary of table output of sPC_n_ (**E**). Bases positive (> 0.05) and negative (< −0.05) in PC3 are shown with sequences of several samples that are indicated in panels **B** and **C** by green or magenta asterisks.

In contrast, each axis of PCA would regards specific connections, presenting structure easier to understand and less distortions. The 1st and 2nd axes of PCA separated samples into four major clusters, each of which is related to the references (Fig 2B), finding bases common to the clusters. The samples were further separated or connected in the lower axes that regarded to other connections of the network structure of species. For example, a cluster found in PC1 and PC2 (Fig 2B, bottom center) was further separated into three groups that include *Bacillus, Mycobacterium*, and *Shinella*, in PC3 (Fig 2C). Hence, *Bacillus* and *Shinella* were similar in part, whereas they were different in other parts; they were gathered and separated accordingly in the corresponding axes of PC for samples. The clusters of samples had different bases at specific positions, dispersed to entire range of the sequence, as obvious in PC for bases (Fig 2D and 2E); bases that presented higher magnitude had more contributions to separate samples.

The relationship of PC for samples and bases appears in Figs 2C (*y*-axis) and 2D, both of which present the PC3 axis. In Fig 2D, positive bases (green) contributed to pulling up *Shinella* in Fig 2C, and negative bases (magenta) contributed to pulling down *Bacillus*; i.e., the green bases tended to appear in *Shinella* samples, while the magenta bases tended to appear in *Bacillus* samples. Bases that had small contributions are shown in gray, and those that had no contribution were omitted. Here, Fig 2D is presented to provide a view of the whole; to specify the bases, scores should be checked on tables. Panel E presents a part of summarized table output, showing PC3 positive and negative bases and some samples including *Shinella* and *Bacillus* sequences (indicated in panels B and C as green and magenta asterisks). Although they are similar as they showed likely PC1 and PC2 values, they differed in several bases in PC3, especially in the sequential indels. As an example of the table output, data are supplied as Supporting Information (S1 Table for bacteria and S2 Table for lion samples).

### Three distinct clusters found in Asiatic lions

Next, let us examine another example of the grouping structure of an engendered species. Nucleotide sequences for major histocompatibility protein (MHC) of Asiatic lion [12], and corresponding sequence of human samples were directly analyzed by PCA (Fig 3). Among the lion’s samples, there were three clusters distinctively separated (Fig 3A). Although differences among the three clusters were comparable to variations among human samples (Fig 3B), differences within the clusters were quite small, and this is contrastive to the well dispersed dots, although a small cluster is separated at the upper right position in human samples.

**Fig 3.**
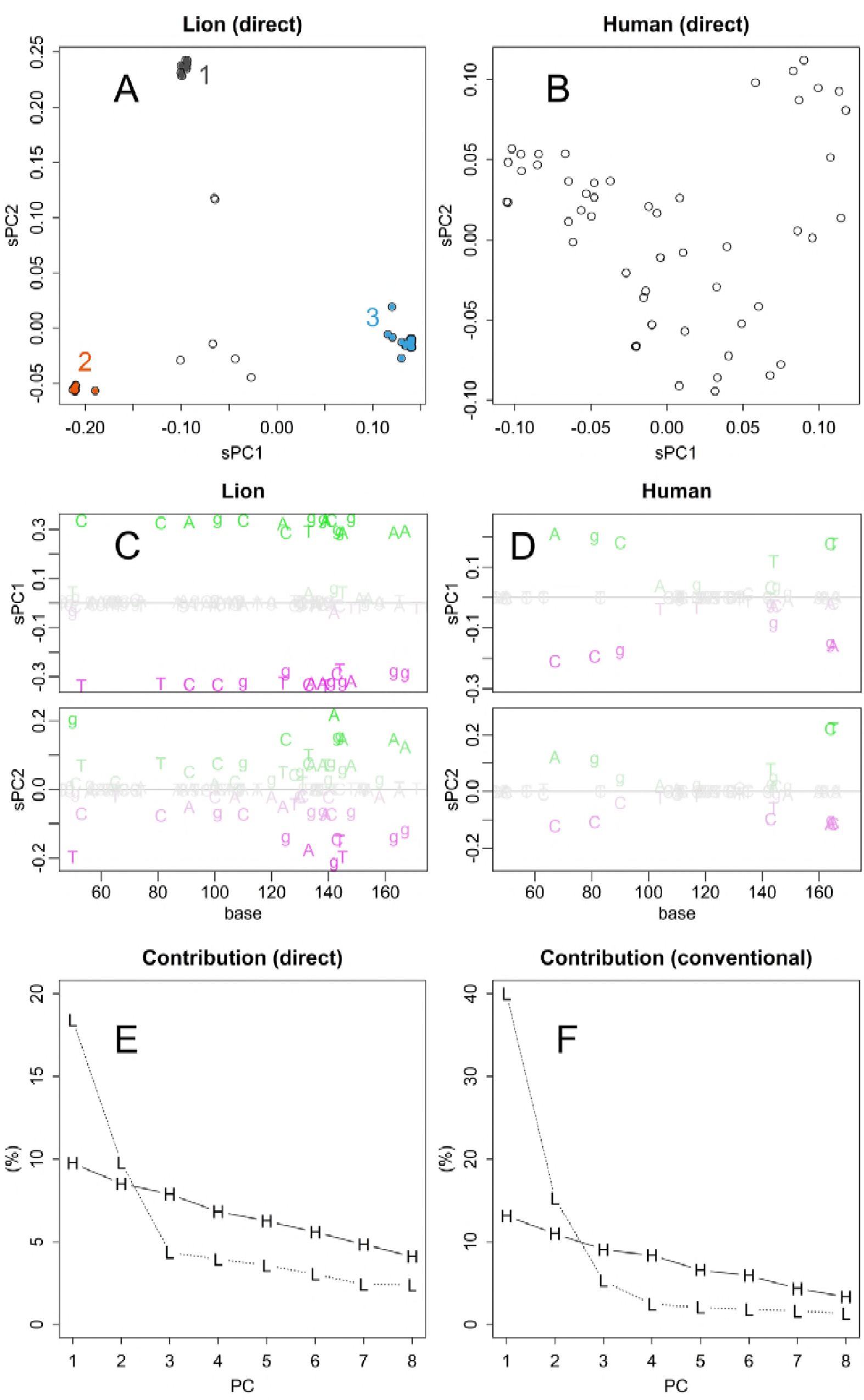
Direct PCA on MHC samples. Scatter plot of PC1 and PC2 for samples of Asiatic lions (**A**). PCA was performed directory to the Nucleotide sequence matrix. Scatter plot for samples of human data (**B**). PC1 and PC2 for nucleotide bases of lion data (**C**). PCs for bases of human data (**D**). Contributions (%) for PC1 to PC8 of PCA applied to sequence matrix directory (**E**) and distance matrix (**F**), respectively.

The different grouping structure between lion and human also appeared in PC for nucleotides. PC1 of lion took extreme and even values (Fig 3C). This shows that each of the tree groups have strong propensity to specific bases at defined position. PC1 for lion presents the difference among the groups, and it contributed 20% of the total differences (Fig 3E). On the other hand, bases took various values both in PC1 and 2, and the number of bases presented is much less in human samples (Fig 3D). Bases differs along larger number of underlying directions; hence contribution of PC1 and 2 were fewer than those of lion and decreased gently (Fig 3E).

Bases that were common to all the samples could be identified, as they had null PC scores in all the axes. Moreover, bases that were specific to a group appeared in characteristic axes. For example, the group A in lion samples appeared as bases that were positive in PC2. Therefore, those bases were automatically selected by specifying the threshold; quantile–quantile plots may suggest the value of the threshold (S1 Fig). These analyses can be performed on a spreadsheet software; however, they can also be performed systematically on R (scripts are in the Supporting Information S1 Doc). As an example of an output, aligned lion sequences are presented with the consensus and characteristic bases in the FASTA format (S6 Table).

### Conventional and direct PCA: Resolution of samples

In the conventional method, the analyzed matrix had been summarized by concentrating to information of distance. Therefore, variations in data could be bundled within fewer axes (Fig 3F). However, the summarizing process would mix up information oriented in various directions. Actually, the groups appeared in PC for samples became more dispersed (Fig 4). The only difference between the two methods, conventional and direct PCA, was the subject of the singular value decomposition: distance and sequence matrix, respectively. Therefore, the dispersing was caused by using the distance matrix. This is a critical characteristic in the classification of samples; the dispersed samples hamper the separation and definition of groups. The distances presented were much longer than those of the direct method. This enlargement was partly provoked by the higher contributions as described above. Another reason is differences in source of distances: in the conventional method, distances between samples are indicated, while those between the center and samples are indicated in the direct method.

**Fig 4.**
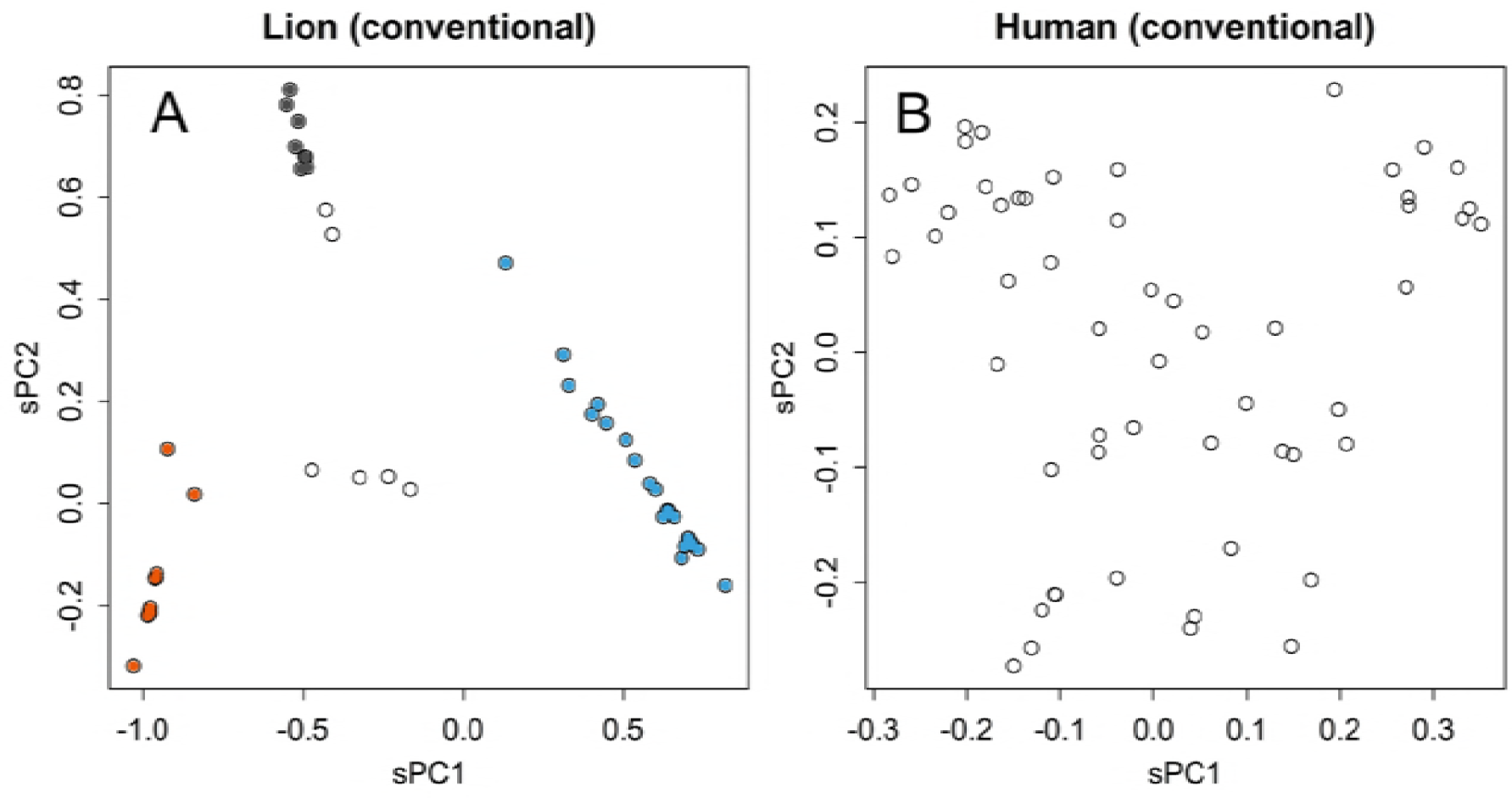
PCA by using distance matrix. Scatter plot of conventional PCA that applied to distance matrix of lion (**A**) and human (**B**) samples.

### Response to outliers

The direct method was robust to outlying sample. A human sample selected by chance contained some unique motifs; the motifs gave long distance to other samples in the distance matrix (S2B Fig). The axes for PC1 and PC2 were mainly determined by this sample in the conventional PCA (Fig 5A); the sample behaved as an outlier, a sample that changes the whole results much.

**Fig 5.**
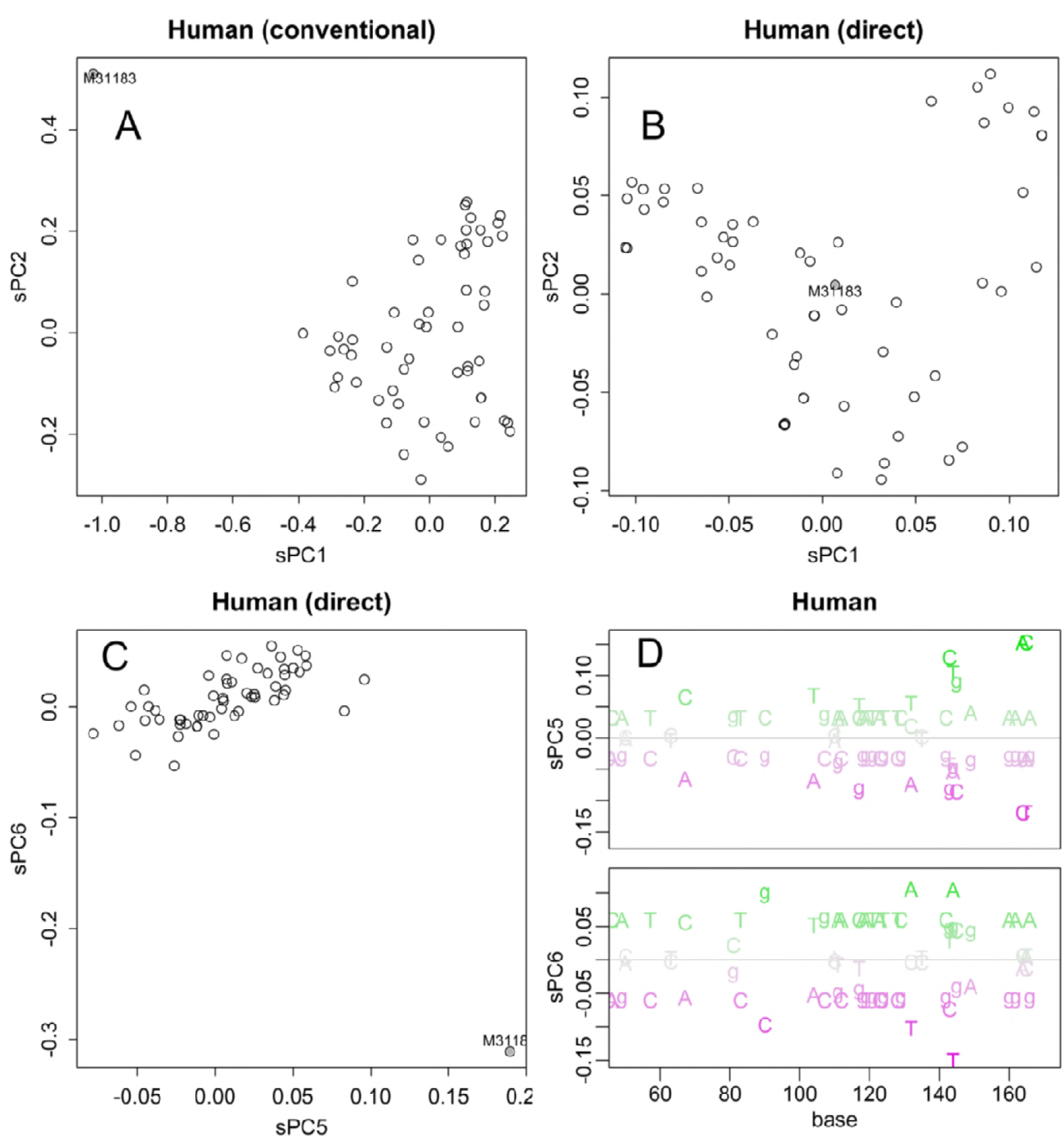
Effect of an outlier sample. Results of PCA of samples that include an outlier. Conventional method applied to the distance matrix (**A**). Direct method applied to the nucleotide matrix (**B**), (**C**). The bases specific to the outlier (magenta, **D**).

By distinguishing the exceptional motifs, the direct method improves robustness to the outlier. In PCA of nucleotide matrix, the sample does not appear as an outlier in both PC1 and PC2 (Figs 5B). Indeed, the whole shape of PCs was almost identical to that calculated without the sample (Fig 3B). The unique motifs gave specific direction that appeared in PC5 and PC6 (Figs 5C and 5D).

### Response to altered alignment

Several methods are available for the alignment of sequences, and the results of the alignment would vary slightly among the methods. However, if the default setting of the parameters was used, they did not alter the results of the direct PCA much (S3–S5 Figs). By reducing the gap penalty to one-third, the average number of gaps per sample increased from 0 to 13 in human data. Such immoderate intervention affected the limited axes; in the case of humans, it altered PC2 and PC3, but not other axes (Figs 6A, 6B, and S6). Conversely, the gathering of multiple clusters was severely altered in the results of the NJ method (Fig 6C). This shows the sensitivity of the NJ method to arbitrary parameters, which would render the results of the analysis dependent on the specific analyst of the data.

**Fig 6.**
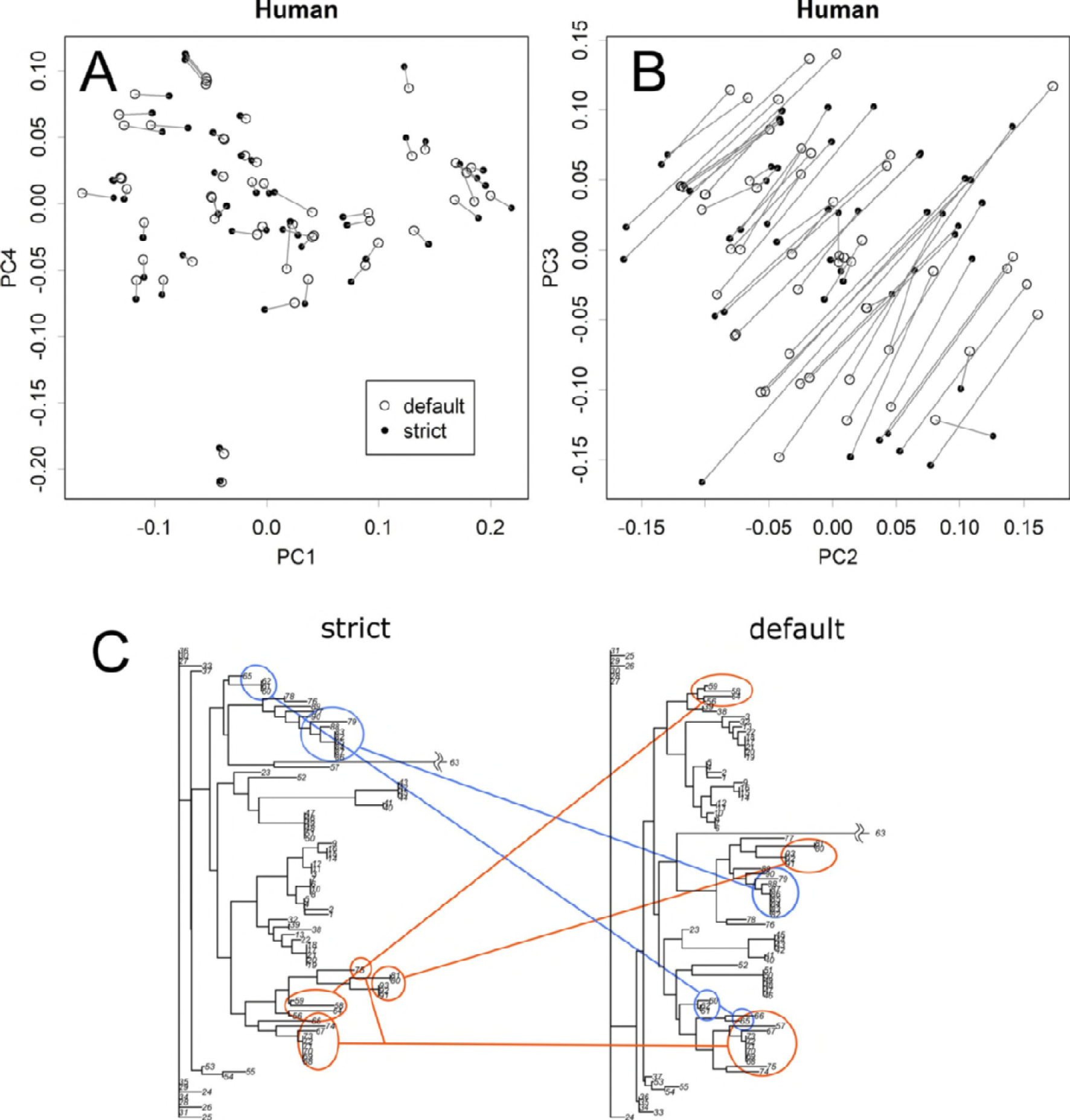
Effect of arbitrary parameters on the alignment. In the alignment process, the gap parameter of the default condition was decreased to obtain strictly aligned sequences. Results of direct PCA (A, B). Identical samples were connected. The effect appeared only in limited axes (B and Fig S7). The connection of clusters was totally altered in the results of the NJ method (C).

### Distance adjustment by JC69 model

The JC69 model will enlarge distances in a matrix; it makes a lattice grid to have spindle shaped skew, and could alter PCA, which finds directions and length solely in the applied matrix. In comparison with Fig 3A, the presented case (Fig 7A) a bit enlarged the differences among clusters; the differences are clear in the distribution of the compensation ratios (Fig 7B). However, as samples in a same cluster would be compensated likely manner, alterations to the relationships were rather limited. Indeed, this did not change much the results of human (S7 Fig).

**Fig 7.**
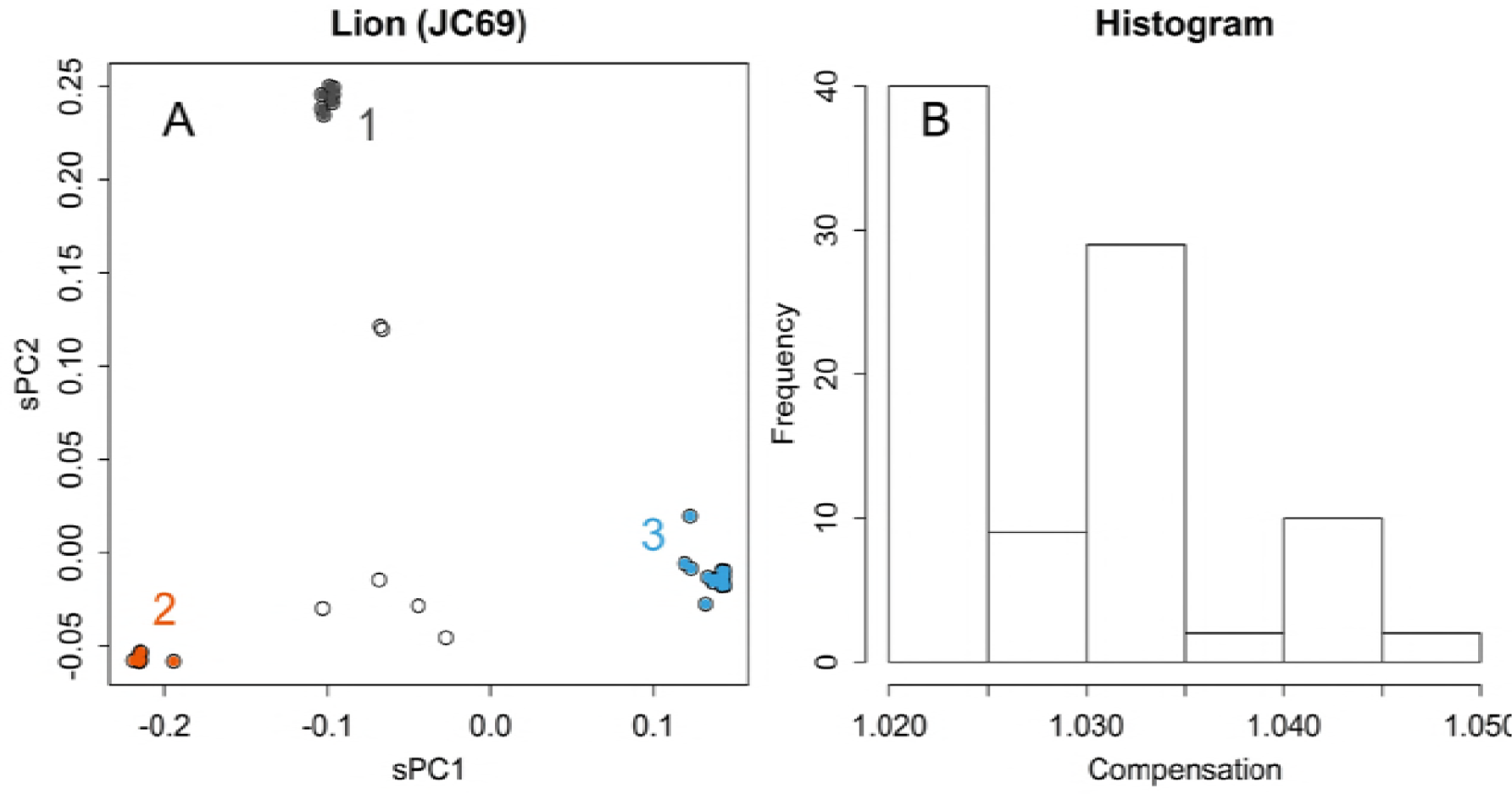
Adjustment of distances. Results of PCA applied to lion samples directory, with an adjustment according to JC69 (**A**). Histogram of ratios of the compensated and original distances (**B**).

## Discussion

Groups that were separated from others in some axes were further separated in other axes (Fig 2B and C). This indicates that they shared certain motifs and differed in others; the shared and different motifs would appear in PC for bases at corresponding axes. This is a typical characteristic of data with a network structure, in which samples are connected by multiple relationships with each other. Direct PCA presents each of the relationships in an axis; the appearance in an axis may be a necessary condition for a real relationship. Hence, similarity in an axis does not always prove genetic relationship; however, if such relationship really exists and if the sequence matrix has information that is sufficient for detection, the relationship will appear in one of the axes. Such multiple relationships will get lumped together and replaced by an average distance in the distance matrix. Hence, conventional PCA will show the relationships in a smaller number of dimensions (Fig 3E and 3F); however, the presented structure will not show the real structure and the resolution will be damaged (Figs 3A and 4A). Moreover, the tree-based methods can give a completely wrong structure; for example, those that appear in Fig 6C are both wrong. By contrast, direct PCA maintains the original information of the sequence matrix, thus presenting the structure of the data in a rotated form.

Asiatic lions were separated into three clusters, and each group has quite small fluctuations (Fig 3A). Differences among the clusters had provoked by specific patterns of sequences (Fig 3C). Most of the differences appeared in the first and second axes of PC, which represented differences of the three clusters (Fig 3E). Unlike the case of human (Fig 3B), Asiatic lions may have propagated from a bottle neck that was formed when they were almost extinct. The distinctively separated clusters may evoke the recently recognized species of giraffe [13]. If the clusters indicate subspecies, each of them has only limited genetic variations. However, there were several samples that showed values in between the clusters (Fig 3A); they seemed to be offspring of hybrids among the clusters, which were not observed in giraffe’s case. If the hybrids are fertile, although the clusters seemed to be isolated with each other, inter cluster mating could retrieve certain magnitude of variety. Such clear separation of sample groups should not be expected in conventional PCA (Fig 4A), and the human samples also became more dispersed (Fig 4B). The decreased resolution occurred because of the conventional method observed in the distance matrix, which mixed up various directions of differences. In this way, the results obtained by direct PCA cannot be negated by those obtained via other methods, which can introduce errors and distortions. It should be noted that direct PCA just rotates the matrix of sequence data; hence, the original structure of the data is maintained.

For identifying unknown samples, both classification of a set of references and relegating the sample into appropriate class are required. In comparison with hierarchical clustering methods, PCA is superior for this purpose in the following points. First, it can separate the classification of references and relegation of a sample; these could be separately performed, as finding of the unitary matrix and applying it to a sample as a multiple regression analysis [11]. Not only reducing calculations, this can protect the system from effect of outliers (Fig 5A) or parameters (Fig 6). Second, the presentation would become easier to understand, especially when number of samples are large. Not limited in the bacterial case shown in Fig 2A, observing the whole structure of NJ, shown within a single dimension, will become rather difficult when numbers of samples are large (for example, that for lion and human’s cases are in Supplementary data as S2 Fig). As PCA does not mix up the dimensions, each of the presentations will become simpler. This characteristic is advantageous in classification in two ways: better resolution (Figs 3A and 4A) and the specification of the base that allowed their separation (Fig 3C). Moreover, the low reproducibility observed indicates a disadvantage in automatic classifications using an inappropriate model (Fig 6C).

Superiority of the direct PCA than the conventional PCA became clear in separation of samples and robustness (Figs 3–5). These base on intactness of information recorded in multivariate data. Additionally, information of base will supply useful cues to understand relationship of sample differences and function of the sequence; this will become clearly apparent in analyzing functional motifs of amino acid sequences.

It should be noted that the robust character of the method may connect to insensitivity to differences that are potentially important. Actually, PCA is a method that observe a matrix from certain direction, and differences among other directions will be ignored. The main purpose of the conventional PCA is to summarize most of differences within the first two axes [9]. The direct PCA to sequence matrix may not be a good tool for this purpose; actually, the contribution of the first two axes were much higher in the conventional method (Figs 3E and 3F). Rather, the direct method is preferable when observing toward lower PC axes, both for samples and nucleotides, to find magnitude of differences and how. Of course, such summarization, i.e., the presentation of a structure using smaller number dimensions, will inevitably introduce distortions in the presented structure.

In the subjects studied, sites that contributed highly to the separation of samples appeared scattered throughout the sequences, rather than gathered to form sequential motifs (Figs 2D, 3C, 3D, and 5D). The characteristics of the sites would appear in each of the axes of PC for samples; which samples are separated and how this occurs is indicated on the axes. The sites may have contributed to the separation, with other sites with similar scores.

The compensation for distance by JK69 did not alter much to the results (Figs 7 and S7). This is partly because of the magnitude of distance, which was less than 0.01 in 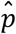 in the presented data. However, this may not be very short in comparing homologous sequences, such as found from database searching. Other more sophisticated Marcov models [1, 2] may be applicable to **C**, however, effects would be rather limited. Additionally, although Markov chain models presumes vertical evolution: i.e., changing each base of a sequence by chance, diversity of sexually reproducing organisms are made by exchanging chromosomes, and those of bacteria are accelerated by horizontal gene transfer. Markov chain models could conflict to those extensive changes.

Since bases may have different weights in physical meanings; for example, the third base of a codon should be lighter than the first, because of the redundancy of the codon. Therefore, **C** could be scaled to equalize weights of positions, by estimating the weight from standard deviations of each column, for example. However, it should be noted that sometimes technical errors may occur in sequencing process. This may give a unique motif among samples; scaling by standard deviation will enlarge artefacts caused by such errors.

### Experimental procedures

Information for MHC1 of 92 Asiatic lions [12], which lives only in the Gir forest sanctuary of Gujarat State in India, had been retrieved from DNA Data Bank of Japan (DDBJ) database [14]. As those lion sequences were not full-length, corresponding region in human gene, HMC1, sequences were also analyzed for comparison; as the human database seemed to have a bias in nationality, 93 were selected out from 217 registered samples, avoiding repetition of contributors of data. One of the samples, accession ID of M31183, differed from others much (Figs 5A, 5D, and S2B); it was selected by chance, and used as an example of outlier.

Nucleotide sequences for bacteria 16S ribosomal DNA were obtained in a student training course held in our department using clone library method or colony direct PCR. Environmental DNA were isolated from active sludge taken from Rinkai sewage plant Akita using ISOIL (NIPPON GENE, Japan) according to the manufactures instruction as instructed in its accompanying manual. 16S ribosomal DNA amplified by polymerase chain reaction using eubacteria specific primer set [13]. And a TOPO TA Cloning Kit (Invitorogen, USA) was used to prepare a genomic library of the PCR amplified genes. Nucleotide sequence of a single colony was determined in Biotechnology Center of Akita Prefectural University, and the sequences were determined by Sangar method [15]. The 16S ribosomal DNA sequence used to analysis were registered under DDBJ [14] accession No. LC279073 to LC279194.

The nucleotide sequences were aligned by using MUSCLE [16]. Other methods (ClustalW [17], MAFFT [18], and DECHIPER [19]) were used to identify the effect of the alignment methodologies. Default parameters used for alignment were applied unless otherwise specified. For the strict and default conditions of DECHIPER, gapOpening was –5 and –17, and gapExtension was –1 and –1.5, respectively. The sequences that were aligned using MUSCLE with accession IDs are presented as Supporting Information (S3–S5 Tables).

### Digitizing the nucleotide sequences

Calculations were performed in R [20]. All the required scripts and sample data are in the supplement data. Nucleotide of length l was digitized by using a boolean vector x of 0 or 1 with length of 5l, connecting information for each base and gap. Then the vectors were piled to make a sequence matrix X. For example, that contains two samples having TAGC and GAGC is,

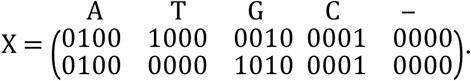

This matrix possesses mutual compatibility with the nucleotide sequence matrix. Any base position consists of the five columns, and the degree of freedom for the five columns is one. As each base is identical, length of differences was estimated in Euclidean length, and then scaled by the number of bases [11].

PCA rotates a matrix to fit the directions of differences to diagonal axes [21]. To do this, center of the rotation have to be defined in prior to the rotation. Mean center of the matrix is a preferable choice, but it could be found from specific sets of reference samples [11]. Samples apart from the center would gain larger moment, and so would have larger influence in finding directions of the differences. The average center is found as means of each columns of the matrix; hence, in the presented example X, it is,

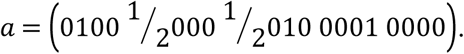

The X could be centered as c = x – a at any rows to give matrix C. The C records differences of each sample from the average sequence at each position of the bases.

### Distance

In direct PCA, differences among samples were estimated by using Euclidean length, as 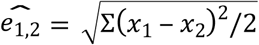, where x1 and x2 are elements of any combination of vectors x1 and x2, scaling the double-counted substitutions at corresponding two nucleotides. This is further scaled by the number of bases, l, after estimating the principal components. It is noteworthy that the scaled component (see 2.5) presents square root of the conventional substitution rate, 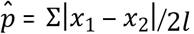.

### Adjusting by a Markov chain model

Distance from the center, which is recorded in C, was optionally adjusted the rate of parallel or back substitutions by a Markov chain model, JC69, which estimate a same frequency of changing for any bases and any positions [22]. For this, scale of c was adjusted to fit the evolutionary distance, by multiplying a compensation ratio. Raw distance of each sample from the average sequence was found as 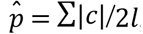, where c is elements of c. Corresponding evolutionary distance was estimated as 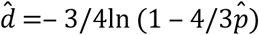. Then each item of C was adjusted as 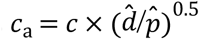, where 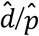 is the compensation ratio uniquely found for a sample.

### PCA

The C or its adjusted version Ca, both of which was a centered nucleotide sequence matrix, was directly subjected for singular value decomposition, C = U∑V*, producing two unitary matrixes U and V, and a rectangular diagonal matrix ∑ that records singular values. The principal components for samples were estimated as PC_s_ = U∑ = CV, and for nucleotides as PC_n_= V∑ = C * U; note that those are rotation of C, without altering relationships among elements. Then they were scaled as 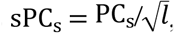 and 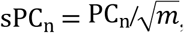, where m is the number of samples, to conform to the mean Euclidean distance [11].

To comparison of methodologies, here the conventional PCA were applied to a distance matrix that uses the same system of distance: not on 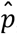, but *ê*. The distance matrix is centered and applied to singular value decomposition. Then components for samples were estimated and scaled for the length of nucleotides; the resulted values were further adjusted for double recorded in the matrix (as it comprises both upper and lower triangles) as dividing by 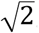.

### Neighbor joining

The distance matrix *ê* was scaled by the same way of the conventional PCA, and further processed by nj function of ape package [23] of R [20].

